# Outer Membrane Vesicle Mediated Multidrug Resistance Gene Transfer in *Avibacterium Paragallinarum*

**DOI:** 10.1101/2022.04.13.488265

**Authors:** Jie Xu, Chen Mei, Yan Zhi, Zhi-xuan Liang, Xue Zhang, Hong-jun Wang

## Abstract

Infectious coryza is an acute upper respiratory tract infectious disease caused by *Avibacterium paragallinarum*, which can cause growth retardation and egg production decline of bred chickens, and bring great economic losses to poultry industry. *A. Paragallinarum* is a Gram-negative bacterium and can release outer membrane vesicles (OMVs). In this study, a comparative genomic analysis of *A. Paragallinarum* isolate P4chr1 and its OMVs were carried out, and the ability to transfer antibiotic resistance genes via the OMVs was studied. The sequencing and data analysis showed that genome size of *A. paragallinarum* P4chr1 is about 2.77 Mb and it has a 25 kb tolerance island covering 6 types of antibiotics and 11 resistance genes. The genome size of its OMVs is about 2.69 Mb, covering 97% genome length and almost all gene sequences of P4chr1. When purified and DNase treated *A. paragallinarum* P4chr1 OMVs were co-cultured with antibiotic sensitive *A. paragallinarum* Modesto strain on an antibiotic containing plate, the colonies grown on the plate were detected corresponding antibiotic resistance gene (ARG). However, antimicrobial susceptibility test exposed that drug resistance genes delivered by OMVs were not persistent, they only existed temporarily on the antibiotic plates. The antibiotic resistance and ARGs disappeared at second bacterial passage. Overall, this study is the first report to compare genomic characteristics of OMVs with its parent *A. paragallinarum* strain, and to study *A. paragallinarum* ARG transfer via OMVs. This work has provided useful data for further study on the issue of non-plasmid ARG transfer mediated by *A. paragallinarum* OMVs.

## NTRODUCTION

Infectious coryza is an acute upper respiratory disease of chickens caused by *A. paragallinarum*, a Gram-negative bacterium of the genus *Avibacterium* within *Pasteurellaceae*. The disease occurs worldwide and leads to serious economic losses in chicken industry due to retardant growth of broilers and reduced egg production in layers [1, 2]. To control the disease, medication is one of the main strategies, so the reatment with antibiotics is optional in many countries. Several studies have reported increased outbreaks and detected antibiotic-resistance genes (ARGs) in recent years [3–5].

Increased appearance and spread of multidrug-resistant bacterial pathogens is one of the major threats to public health [6] and food animal health worldwide. Because along with the emergence and development of bacteria on earth, their resistance to environmental pressures keeps growing. Wide application of antibiotics in human and food animal industry lays such a pressure to the life of bacteria. In return, more and more bacteria gradually generate resistance against more and more drugs. The speed of occurrence of drug-resistant bacteria is found surpassing the pace of new antibiotics created [7]. Meanwhile bacterial drug-resistance could transfer from food animal to human through accumulated drug residue in meat and egg products.

It has been recognized that tackle the infection caused by Gram-negative bacteria is more difficult than Gram-positive bacteria as they have two-layers of sophisticated cell membranes [8–9]. In order to adapt to the adverse environmental conditions, Gram-negative bacteria have evolved a globular bi-layered vesicle (diameter 50–500 nm), named outer membrane vesicles (OMVs), which are derived through blebbing and pinching-off from bacterial outer membrane without destroying it [6, 10]. OMVs contain many of the same components as the outer membrane of the bacterial cell, such as lipopolysaccharide, phospholipids, membrane proteins, and peptidoglycan components [6, 10]. The lumen of the vesicles contains periplasmic proteins, cytosolic components, and nucleic acids [11]. OMVs also carry DNA and RNA on their surface, which can be removed by OMV treatment with DNase and RNase, whereas luminal DNA and RNA are nor affected by the treatment [12].

Recent studies have revealed a new mechanism for antibiotic susceptible bacteria to gain ARGs from ARG donor bacteria (in same species or different species) using OMVs as vehicles, other than the three traditional routes: natural transformation, transduction or conjugation by bacterial cells [6]. Although these known mechanisms contribute to the gene flow within bacteria, they have restrictions such as limited genetic load, host specificity, and the kind of genetic material to be transferred [6].

So far, function and roles of OMVs on ARG and virulent gene transfer and dissemination have been demonstrated in numerous Gram-negative bacteria, such as *Acinetobacter baumannii, Escherichia coli* and *Pseudomonas aeruginosa*, resulting in the expression of obtained genes [13–17]. The work on *A. paragallinarum* has not been reported.

Recently, we sequenced whole genome of a newly isolated *A. paragallinarum* strain and identified several ARGs in it that intrigued us to investigate ARG transfer by this isolate. As *A. paragallinarum* is a pathogen of respiratory disease, research on ARG transfer and dispersing through *A. paragallinarum* OMVs may help discovery of its correlation and mechanism related to respiratory tract invading bacteria.

In this study, we present for the first time the comparative genomics analysis between a multi-drug resistant strain of *A. paragallinarum* P4chr1 and its OMVs, based on genome sequencing information. In addition, also the first we demonstrated *A. paragallinarum* P4chr1 OMVs mediated aminoglycoside antibiotic genes transfer to a drug susceptible strain by horizontal gene transfer (HGT) experiment.

## MATERIALS AND METHODS

### Bacterial Strains

*A. Paragallinarum* P4chr1 was isolated from infraorbital sinus sample of a diseased bird from a chicken farm in China, in 2021 and it was previously identified to be serovar A. 16S rRNA gene sequencing and biochemical analysis were used to identify bacterial species. *A. paragallinarum* Modesto is serovar C reference strain preserved in the laboratory. (GenBank accession number CP086713.1).

### Isolation and Purification of OMVs

The isolation and purification steps of omv were modified on the basis of previous literature [18]. *A. paragallinarum* strain P4chr1 was inoculated in tryptic soy broth (TSB) containing 10% (v/v) chicken serum and 0.0025% (w/v) nicotinamide adenine dinucleotide (NAD), and cultured at 37 ° at 180rpm for 15 hours [19]. The next day, the culture was inoculated into 2L TSB at a ratio of 1:50, and cultured under the same conditions for 12-16h to make its OD600 reach 1.0. Then, the cultured bacteria liquid was centrifuged at 4 ° C 7500 × g for 15 min, the discarded bacteria were precipitated, and the supernatant was collected. The collected supernatant was filtered by Stericup filter (Millipore Corporation, United States) with pore diameter of 0.45 μm to remove the bacteria floating in the liquid. The filtered supernatant was centrifuged at 150,000 × g, 4 °C for 3 h (SW40 Ti rotor, Beckman-Coulter, Germany), the supernatant was discarded, the precipitate was collected, and resuspended with 30 mL of 0.05 mol/L Tris-HCl buffer (pH 8.0). The same process was repeated again, and the pellet was suspended with 5 mL phosphatebuffered saline (PBS) buffer to obtain crude OMVs. The crudely extracted OMVs were centrifuged at 75,000 × g for 1 h at 4 °C, the supernatant was discarded, the pellet was collected and resuspended in 5 mL of 50 mM HEPES-150 mM NaCl solution, and then centrifuged by density gradient centrifugation, add 2 mL series gradient concentration of OptiPrep-iodixanol (Fresenius kabi Norge AS, Oslo, Norway) gradient cell separation solution to the centrifuge tube, the concentration from bottom to top is 45%, 40%, 35%, 30%, 25% and 20% (w/v), the top layer adds 2 mL of crude OMVs resuspended, centrifuged at 150,000 × g overnight at 4 °C. After the completion of density gradient centrifugation, the separation liquid of each layer was collected from top to bottom, each fraction was subjected to SDS-PAGE gel electrophoresis, and stained with Coomassie brilliant blue to observe the distribution of proteins in each layer. The protein-containing density layers were combined and centrifuged at 150,000 × g, 4 °C for 1 h to remove the gradient solution, the supernatant was removed, the pellet was collected and suspended in 50 mM HEPES-150 mM NaCl solution, and then Centrifuge, remove the supernatant and resuspend, and repeat this centrifugation 3 times to fully remove the gradient. Finally, purified OMVs were obtained by resuspending the collected pellet in 2 mL of 50 mM HEPES-150 mM NaCl solution.

### DNA preparation and sequencing of OMVs

DNA was extracted as previously discribed [18]. Since the nucleic acid sequence contained in OMVs is a small molecule sequence, and nanopore third-generation sequencing characterized by long-read sequencing, the genome of OMVs was sequenced through second-generation sequencing.

### Transmission Electron Microscopy

The vesicle suspension was fixed first with cold 2.5% (v/v) glutaraldehyde at 4°C for 2 hours, followed by 1% (w/v) osmium tetroxide for 1 hour at 4°C. After washing with deionized water, the immobilized vesicles were dropped onto a 200-mesh grid and imaged using a Philips CM 100 transmission electron microscope (TEM) at 80 kV.

### Whole genome sequence (WGS) and bioinformatic analysis of *A. paragallinarum* P4chr1

Bacterial genomic DNA was extracted using the Invitrogen dna Mini Kit. (Thermo Fisher Scientific, the United States). *A. paragallinarum* P4chr1 was subjected to WGS using a combination of Nanopore PromethION (Oxford Nanopore Technologies, Beijing, China) and illumina novaseq6000 (Genewiz, Beijing, China) platforms. Use Canu v1.5 and Falcon v0.3.0 to perform mixed assembly of the original data, and use the second-generation small fragment data to perform single-base correction (GATK) on the assembly to obtain a high-confidence assembly sequence. Gene prediction was performed using Prodigal software (PROkaryotic Dynamic programming Gene-finding Algorithm), because compared with other softwares, Prodigal has high quality gene structure prediction and better translation initiation site prediction, and give fewer false positives[20]. Prodigal, glimmer and GeneMark HMM are all used for gene prediction of NCBI prokaryotes. Genomes were annotated using the online database RAST (http://rast.theseed.org/FIG/rast.cgi) and results were corrected using the BLASTn database (https://blast.ncbi.nlm.nih.gov/Blast.cgi). ResFinder database was used to detect antimicrobial resistance genes (ARGs) in genome. (https://cge.cbs.dtu.dk/services/). Gene function and metabolic pathways predictions were obtained with the Blast2GO annotation pipeline. The BRIG (BLAST Ring Image Generator) tool was used to draw the circular map of *A. paragallinarum* P4chr1 and compare it with OMVs. Transmembrane domains (TMDs) in the P4chr1 genome were predicted using TMHMM Server v.2.0. Finally, the software NCBI Blast+ was used to compare amino acid sequence of the protein with the data of COG, KEGG, VFDB and other databases to obtain the protein function annotation information.

### Recipient Cell Preparation

*A. paragallinarum* Modesto was grown in 10 ml TSB with the supplement as above at 180 rpm for 15 h at 37°C. The 2% (v/v) culture was then transferred to 500 mL TSB with same supplements and incubated to OD 0.4-0.6. Subsequently, the growth was pelleted by centrifugation at 4,000rpm for 30 min at 4°C, and the supernatant was discarded. The equal volume of pre-cooled 272mM sucrose solution was used to resuspend the precipitate. Then cells were pelleted by centrifugation at 4,000rpm for 30 min. A 1ml of cooled 10% (v/v) glycerol was used to resuspend the precipitate, and then each of 200ul was packed separately.

### OMV Mediated Gene Transfer

Transformation experiment was performed according to a method described previously by Fulsundar et al [12] (Fig 1). The experiments was conducted in triplicate and at three independent times. To prepare the gene transfer incubation mixure, adding 50 μL of recipient cells into 500 μL of Super Optimal broth with Catabolite repression (SOC) medium in each Eppendorf tube. Then 500 μL of purified OMVs with known protein concentration (mg/mL) and DNA content was added, followed by adding 1 μL of 100 μg/μL DNase (final concentration 100 ng/mL) (Thermo Fisher Scientific, CA, United States). The tubes were statically incubated at 37 ° for 4 h. After the incubation, mixture was transferred aseptically to a culture tubes, and incubated further for 2 h with shaking at 150 rpm. Next, adding 2 mL of SOC medium per tube, and continuing the incubation with shaking for additional 21 h. After this 24 h of incubation, the cells were pelleted by centrifugation, and resuspended in 1 mL SOC medium followed by plating the suspensions on TSA plates with antibiotics or without. Four test groups were set in the experiments by plating on TSA with 4 different antibiotics (Table 3). Meanwhile 3 controls were set: Control A was was prepared as same as test groups, but added no DNase in its sample mixture and plated on TSA without any drugs; Control B was prepared as same as test groups and plated on TSA without any drugs; Control C contained only recipient cells and was plated on TSA with the 4 antibiotics as used in the test groups. The plates were incubated at 37°C for 2 days, and then evaluated by counting number of colonies or transformants grown on each plate for every group. The percentage (%) of transformants was determined by dividing the number of colonies / transformants in each test group with the number of colonies / transformants in control A group.

**FIG 1.**
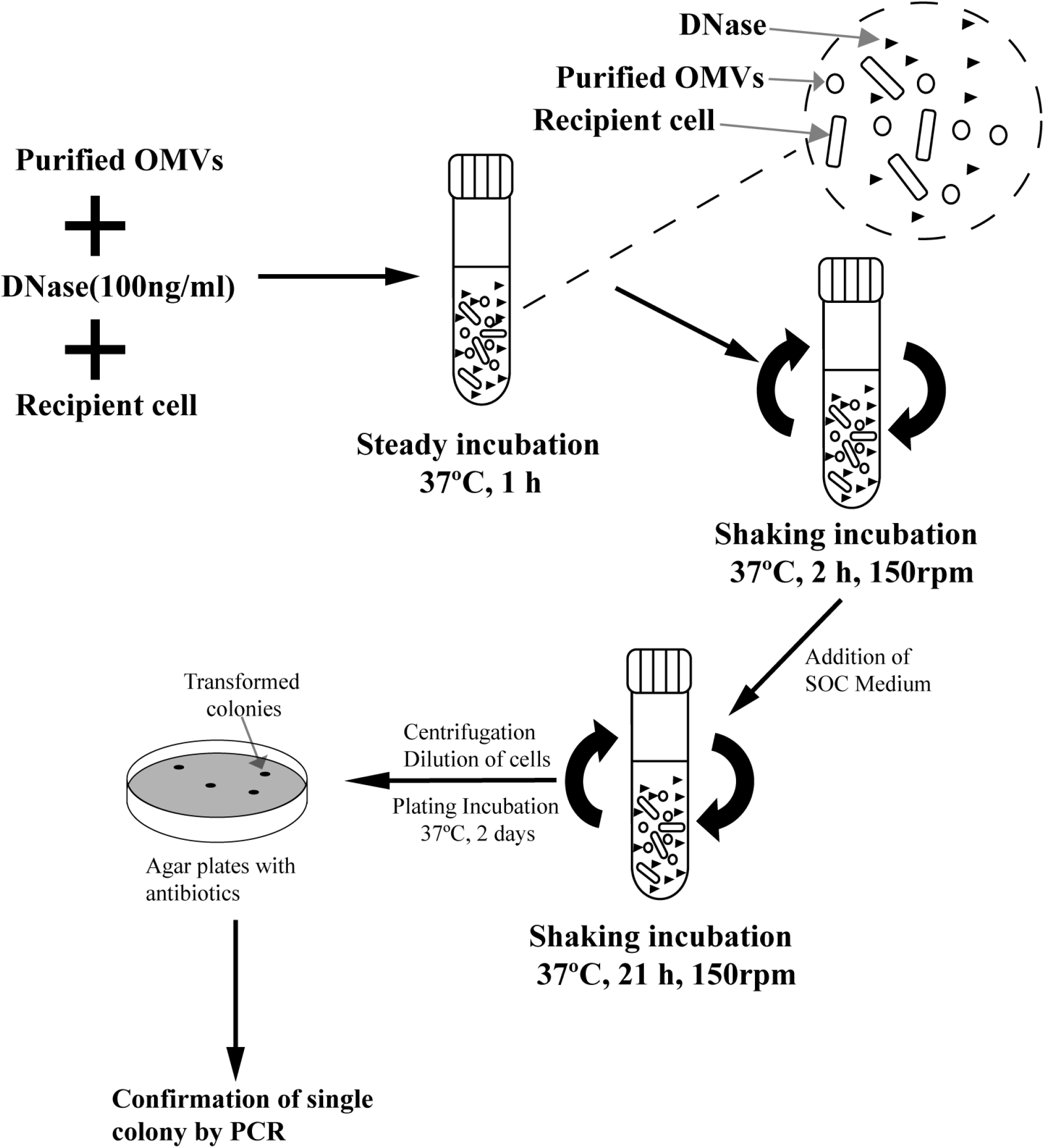
Schematic presentation of the steps involved in vesicle-mediated transfer in *A. paragallinarum* strain [12].

### Confirmation of Gene Transfer by PCR

Based on the genomic information, 4 pairs of antibiotic resistant gene primers were designed and synthesized (Table 1), and PCR verification was performed on a single colony from the resistant plates. The donor bacterium P4chr1 containing DNA with antibiotic resistant gene was used as positive control, and water was used as negative control.

**Table 1.**
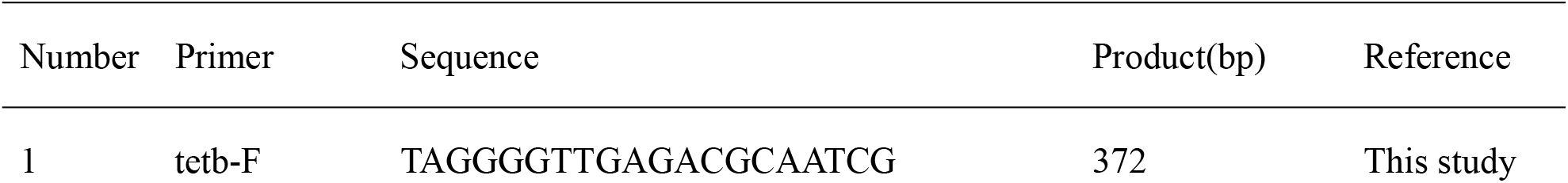
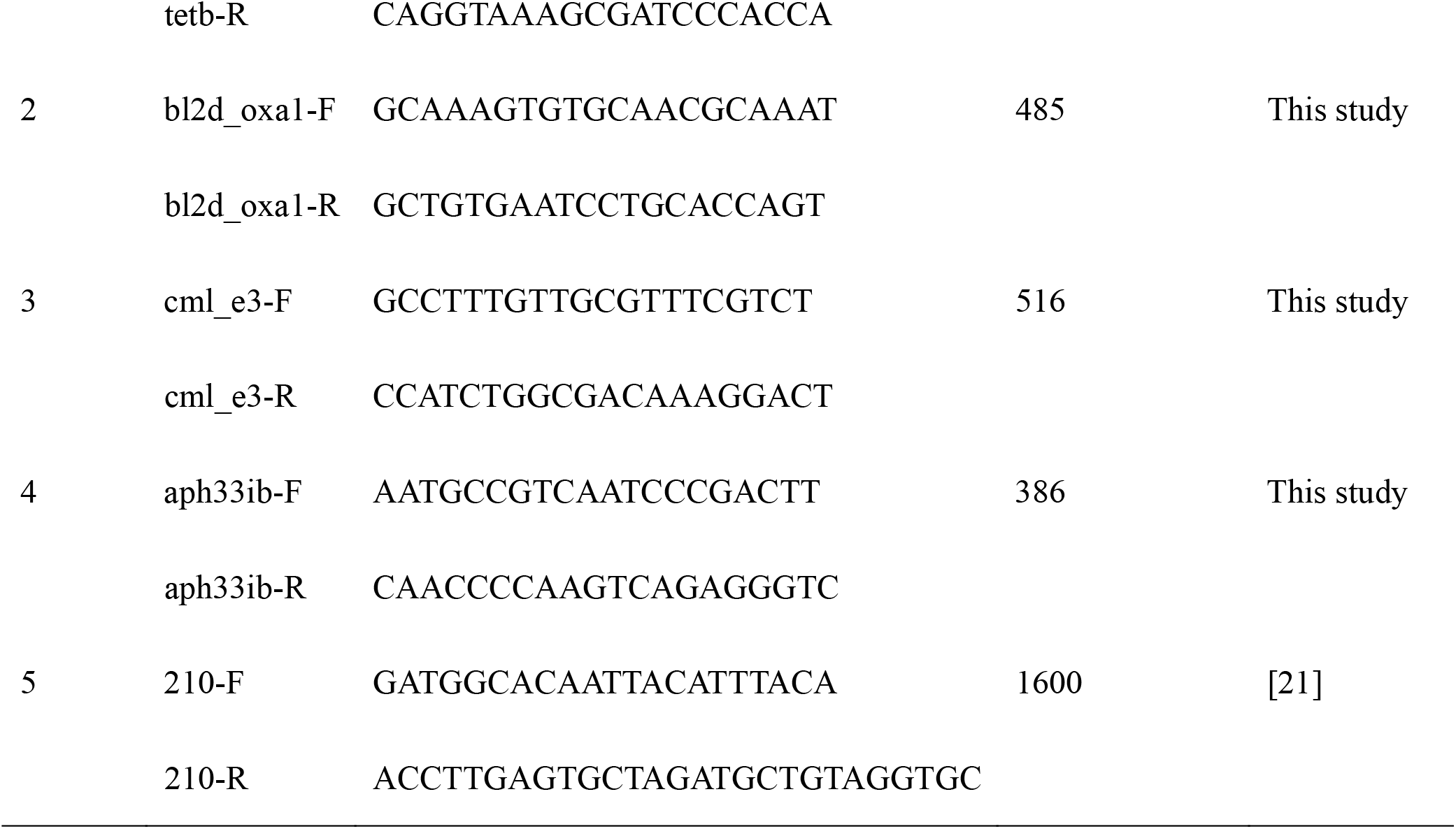
Primer sets used for amplification of the DNA fragment

### Confirmation of Recipient Cell by Serotypeing and PCR-RFLP

The classical HA-HI (hemagglutination-hemagglutination inhibition) test [22]. HA antigens were prepared from TSB cultures seeded with acquired transformants, donor strain P4chr1 (serovar A) and recipient strain Modesto (serovar C). PCR-RFLP (Restriction fragment length polymorphism) analysis [23] were performed on the recipient bacterium, donor bacterium, and colonies on the resistant plates as described above.

### Antimicrobial susceptibility testing

*A. paragallinarum* P4chr1, Modesto and transformants were cultured in TSB with suppliments without antibiotics. Antimicrobial susceptibility testing was performed by a broth microdilution method according to the protocol described by the CLSI (Clinical and Laboratory Standards Institute) [24]. Resulted MICs (Minimum Inhibitory Concentration) data was interpreted according to the recommendations given in CLSI documents VET088 [24] and M100 [25]. *E*.*coil* ATCC 29213 was served as quality control strain.

### Nucleotide sequence accession numbers

The complete genome sequence of the chromosomal DNA P4chr1 has been deposited in GenBank under accession number CP081939, and genome sequence of its OMVs had been deposited in BioSample under accession number SAMN22170838.

## RESULTS

### Ultrastructure of OMVs

The P4chr1 bacterial solution and the extracted OMVs were observed by transmission electron microscope (TEM), and spherical structures were observed around P4chr1 (Fig. 2a), and the same extracted OMVs were similar to this structure (Fig. 2b). It is proved that P4chr1 can secrete OMVs to the environment during growth. The particle size of OMVs is mainly distributed between 30 and 100 nm, and the average particle size is 40 nm.

**FIG 2.**
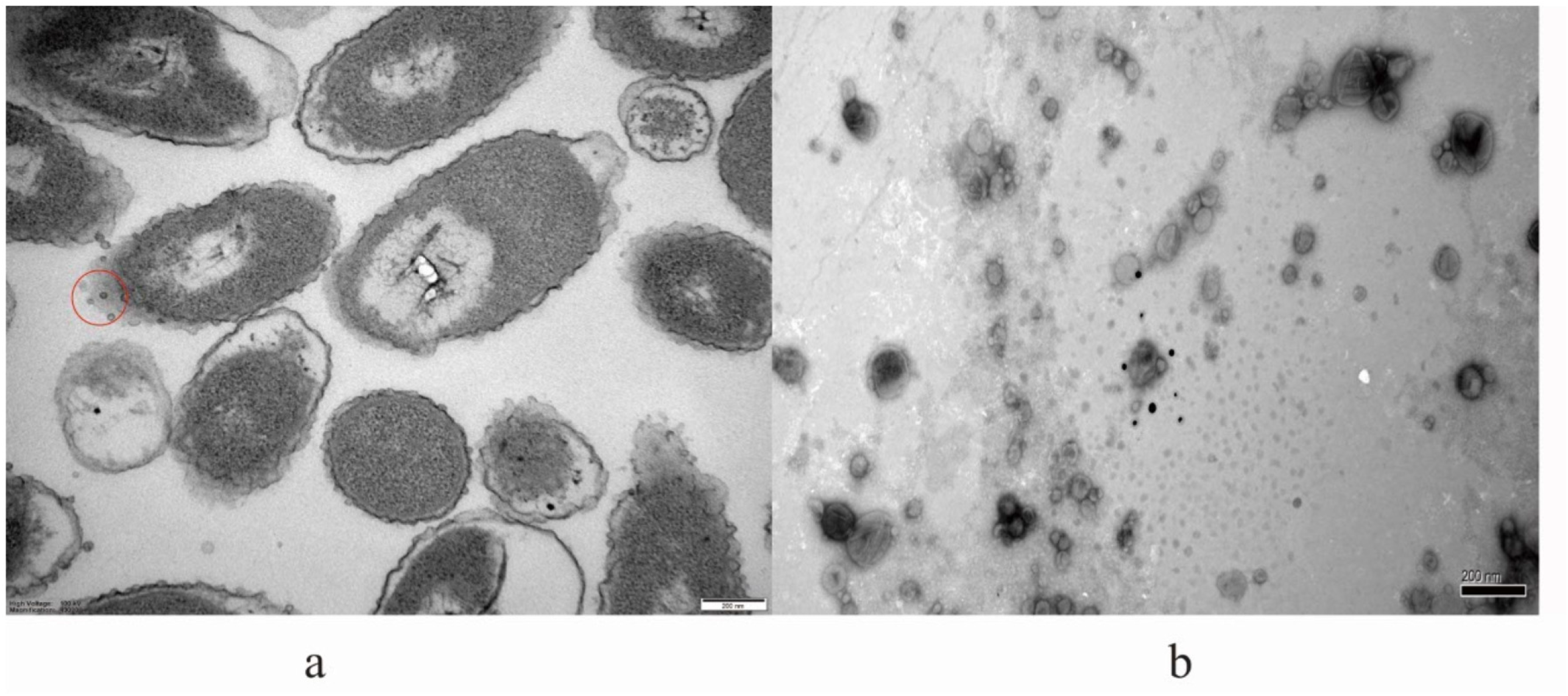
(a) Observation results of *A. paragallinarum* p4chr1 under electron microscope. (b) OMVs released by *A. paragallinarum* field strain p4chr1. Vesicles were purified from broth culture by ultracentrifugation and filtered through a 0.22μm filter. The average diameter of the vesicles was 40 nm. The OMVs were free of bacterial contamination.

### Genome Characterization of *A. paragallinarum* P4chr1

The assembled whole genome sequence revealed that *A. paragallinarum* P4chr1 harboured a circular chromosomal DNA (2,774,989bp) with a 41.01% G + C content, and it did not carry a plasmid. In total, 2,778 protein-encoding genes were predicted, with a coding percentage of 95.92%. The average gene length was 852 bp (Fig 4a). The software Aragorn was used to predict tRNA, and the predicted number was 59. The software RNAmmer was used to predict rRNA, and the predicted number was 15. In addition, the genes were searched against the KEGG, eggNOG, Nr, Nt and Swiss prot databases to annotate the gene description. Among the 2,778 genes predicted in *A. paragallinarum* P4chr1, 2,778 genes were annotated into the COG database, accounting for 92.22% of the predicted genes. According to the COG classification standard, which can be divided into 21 categories. There were 1851 genes being annotated into the KEGG database, accounting for 66.63% of the predicted genes, and these genes can be divided into 38 metabolic pathway types (Figure 3a). Details were in Table 2.

**Table 2.**
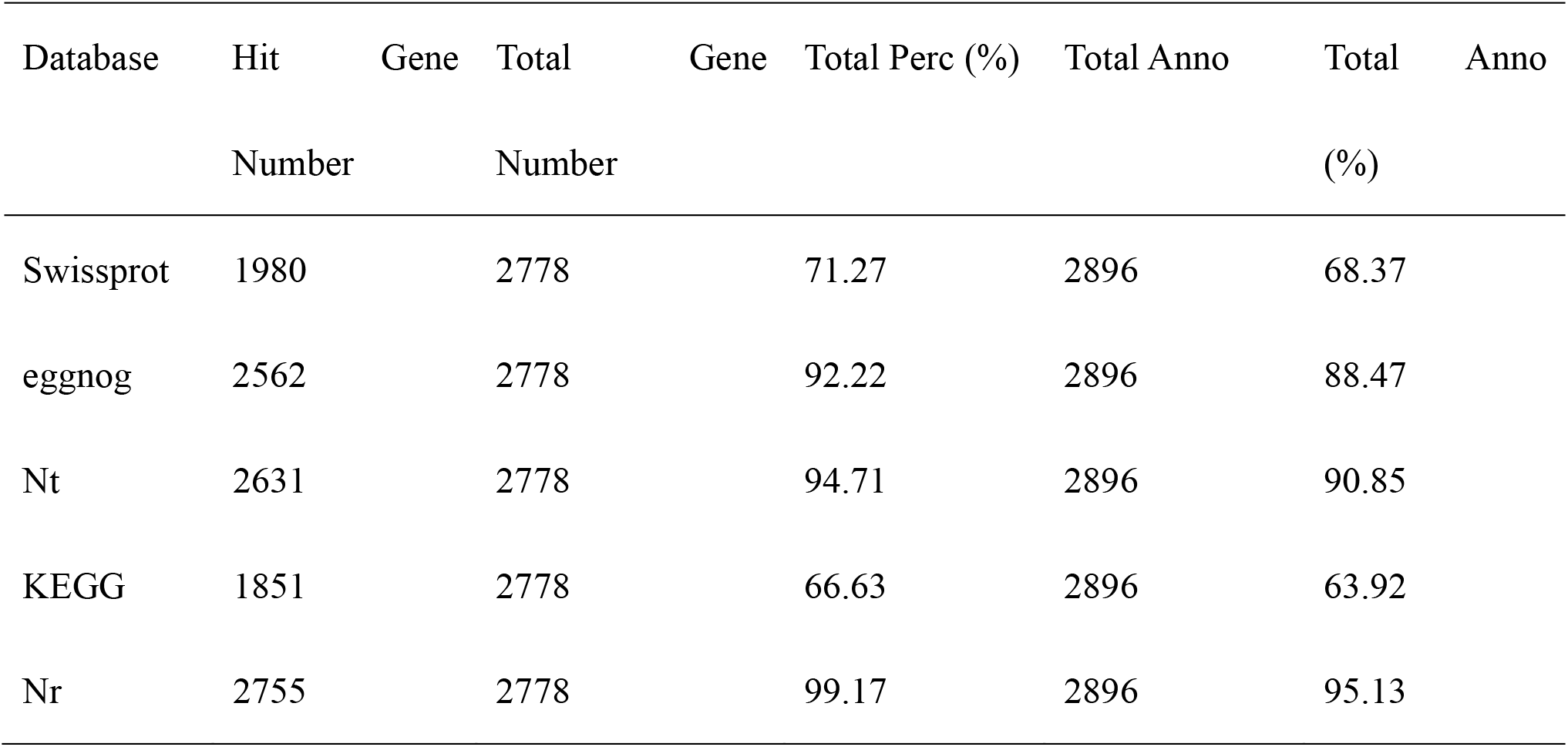
The status of genome of A. paragallinarum P4chr1 annotated in 5

**FIG 3.**
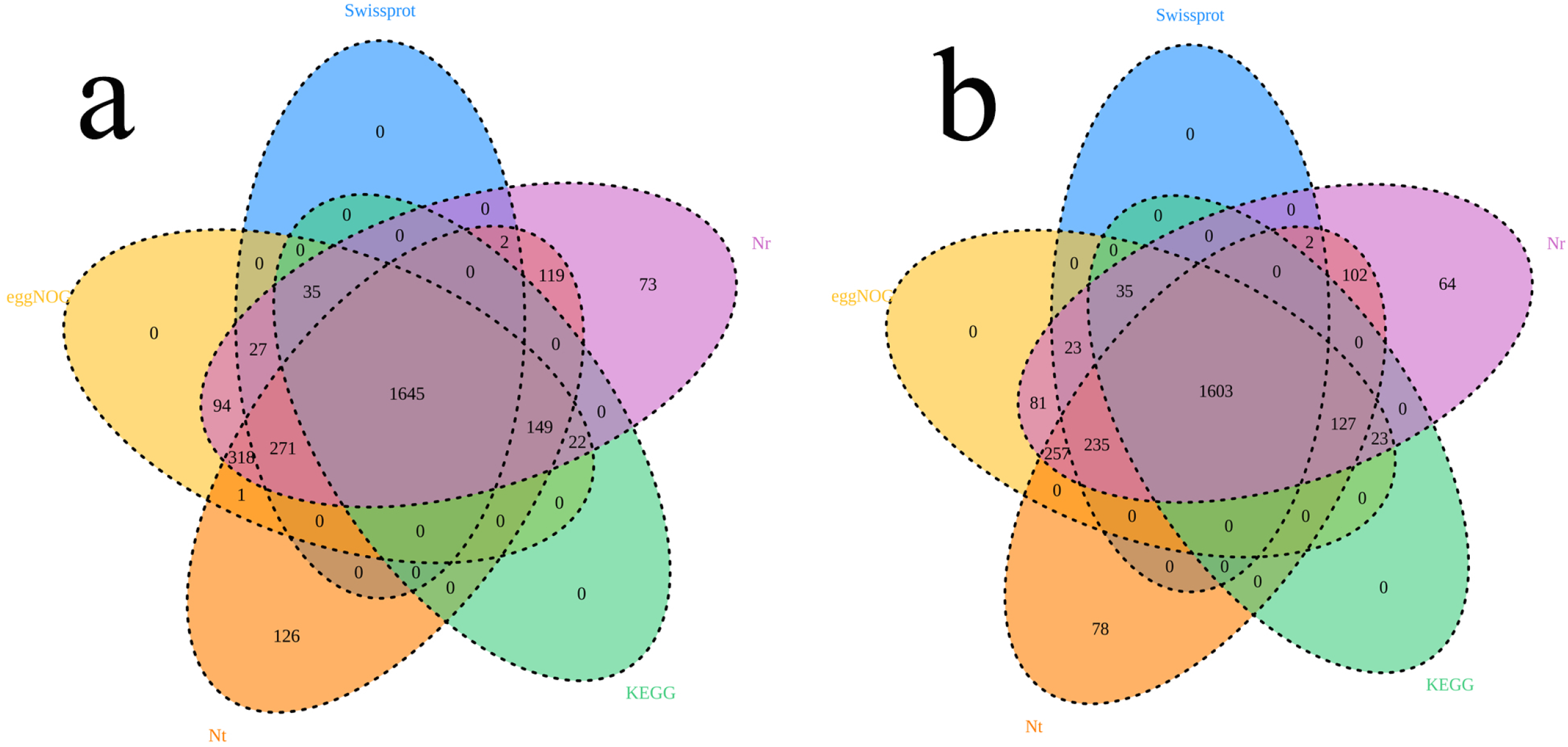
(a) The venn diagram of *A. paragallinarum* P4chr1 genome annotated in each database. (b) The venn diagram of the OMV genome annotated in each database. Different colors represent different databases. Different color circles have overlapping areas that can be annotated by different databases.

**FIG 4.**
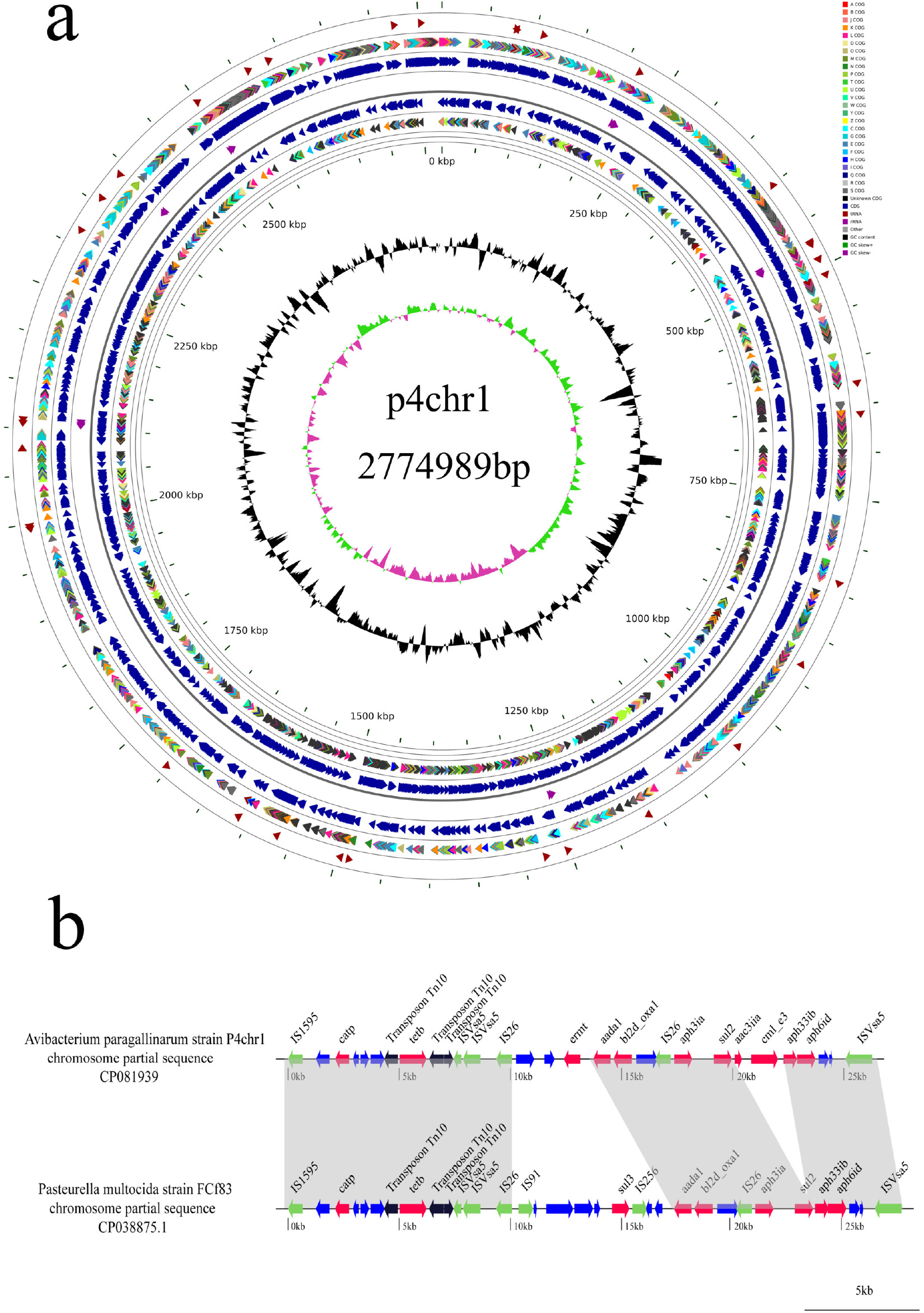
(a) Circular presentation of the map of *A. paragallinarum* P4chr1. From outside to inside: tRNA related genes; COG annotations of forward genes-distinguished by different colors (see instructions in the upper right corner); the position of forward genes; rRNA genes; reverse gene coordinates; reverse gene COG Note: The GC content is based on the average GC. Outwardly protruding means higher than the average, and inwardly protruding means lower than the mean; the innermost circle is the GC skew value, purple means less than 0, and green means greater than 0. (b) Comparison of *A. paragallinarum* P4chr1 with chromosomal corresponding regions of *Pasteurella multocida* strain FCf83 from China. The arrows indicate the extents and directions of transcription of the genes. ORFs with different functions are presented in various colours. The regions with >99% homology between *A. paragallinarum* P4chr1 and the chromosome of *Pasteurella multocida* strain FCf83 are indicated by grey shading.

### Antibiotic Resistance of *A. paragallinarum* P4chr1

The ResFinder results showed that 11 antimicrobial resistance genes of 6 categories of antibiotics [the aminoglycoside resistance gene aph6id, aph3ia, aac3iia, ant2ia and aph33ib, the beta-lactam resistance gene bl2d_oxa1, the MLS (Macrolide, Lincosamide and Streptogramin B) resistance gene ermt, the phenicol resistance gene catp and cml_e3, the sulphonamide resistance gene sul2 and the tetracycline resistance gene tetb] were present in the chromosomal DNA of *A. paragallinarum* P4chr1. The resistance genes in the genome were all concentrated in a 25kb fragment. Sequence comparison revealed that the resistance region of *A. paragallinarum* P4chr1 exhibited high homology to the corresponding region in the chromosomal DNA of *Pasteurella multocida* strain FCf83 (accession no.CP038875.1) from China (Fig 4b).

Antimicrobiol susceptibility testing showed that *A. paragallinarum* P4chr1 was resistant to chloramphenicol, erythromycin, gentamicin, tetracycline, streptomycin and ampicillin, whereas the recipient strain *A. paragallinarum* M was sensitive to these antibiotics.

### Collinearity analysis of the two genomes

The genomic sequence of the OMVs of *A. paragallinarum* strain p4chr1 were composed of 162 contigs for 2,691,804 bp with a 40.92% G + C content. The base pair numbers in the OMVs was 97.00% of that in *A. paragallinarum* p4chr1. The largest contig was 149543 bp, and the minimum contig was 261 bp. In total, 2,568 protein-encoding genes were predicted. The average gene length was 859 bp. The figure 3b showed the OMVs genomes annotated in various databases. The results of the comparative genome circle graphs of *A. paragallinarum* P4chr1 and OMVs showed that the similarity of the two genomes was greater than 90% (Fig 5a).

**FIG 5.**
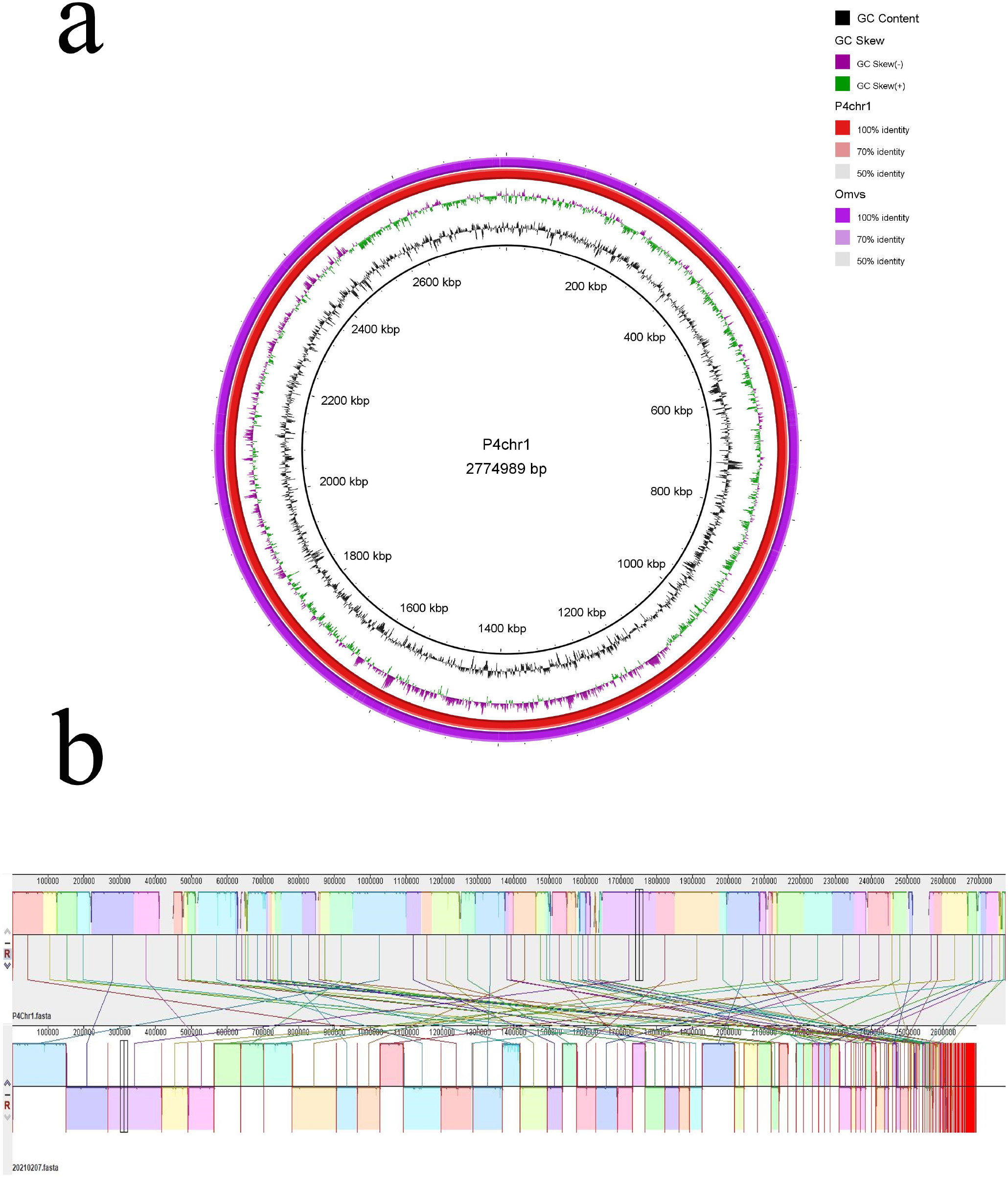
(a) Comparative genomic circle diagram of *A. paragallinarum* P4chr1 and OMVs. The circles show (from outside to inside): OMVs genome sequences, P4chr1 genome sequences, GC skew, GC content and scale in kb. (b) Genome collinearity analysis of *A. paragallinarum* P4chr1 and OMVs. The two parts connected by the line were similar sequences.

The result of collinearity showed that most of the genome fragments of OMVs were found the corresponding parts in the genome of *A. paragallinarum* P4chr1 (Fig 5b). The analysis of whole-genome orthologous clusters was an important step in comparative genomics research. Identifying clusters between orthologous clusters and constructing networks can help us to explain the functions and evolutionary relationships of proteins across multiple species. The result of orthologous cluster analysis of *A. paragallinarum* P4chr1 and OMVs showed that P4chr1 had 2546 clusters, OMVs had 2544 homologous clusters, of which 2541 homologous clusters were shared by P4chr1 and OMVs (Fig 6). These results indicated that the genome of OMVs was derived from *A. paragallinarum* P4chr1, and OMVs had almost all the genome sequence of *A. paragallinarum* P4chr1, including some virulence genes and antibiotic resistance genes.

**FIG 6.**
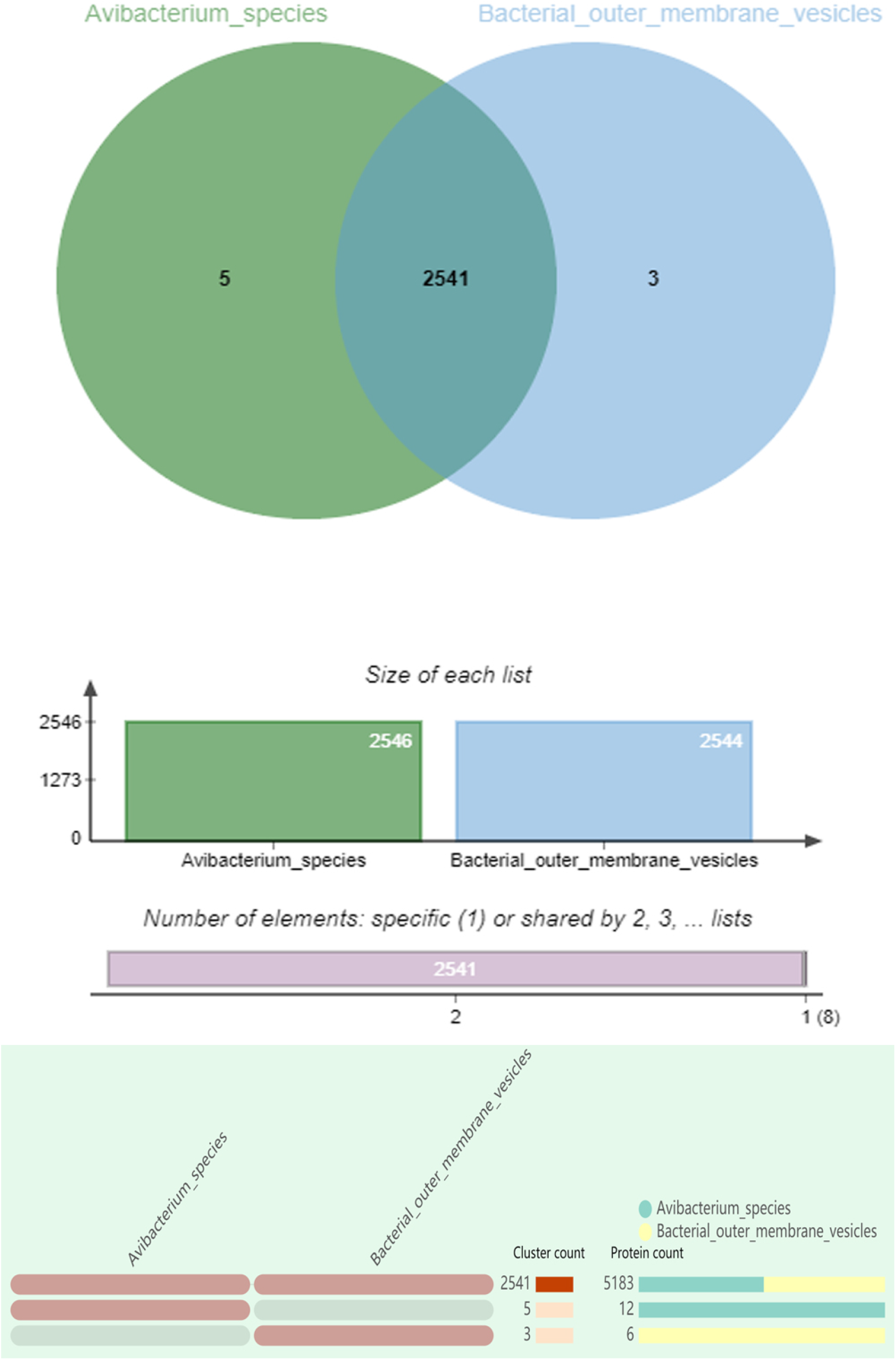
Orthologousclusters analysis of *A. paragallinarum* P4chr1 and OMVs.

### Transfer of Antibiotic resistance genes (ARGs)

Purified OMVs from *A. paragallinarum* P4chr1 were inoculated to TSA and TSB with suppliments, and no bacterial growth was detected after 24 or 48 h of incubation, indicating that the OMVs were free of bacterial contamination.

Following *A. paragallinarum* Modesto was transformed with purified OMVs (Table 3), the transformants produced from resistant plates were tested for ARGs by PCR, and agarose gel showed the corresponding ARG bands (Fig 7).

**Table 3.** OMV-mediated transformant yield obtained after 24 hours incubation with OMVs in TSA agar plates under different treatment conditions

| Treatment<sup>a</sup> | No. of transformants | % of transformants<sup>b</sup> |
| --- | --- | --- |
| Control A | 4.76*10<sup>7</sup> | 100 |
| Control B | 3.54*10<sup>7</sup> | 74.37 |
| Control C | 0 | 0 |
| Test 1<sup>tet</sup> | 13 | 2.73*10<sup>-7</sup> |
| Test 2<sup>amp</sup> | 16 | 4.52*10<sup>-7</sup> |
| Test 3<sup>chl</sup> | 9 | 1.74*10<sup>-7</sup> |
| Test 4<sup>str</sup> | 28 | 5.42*10<sup>-7</sup> |

<sup>a</sup> See Materials and Methods.
<sup>b</sup>Data are the mean values from triplicate experiments.

**FIG 7.**
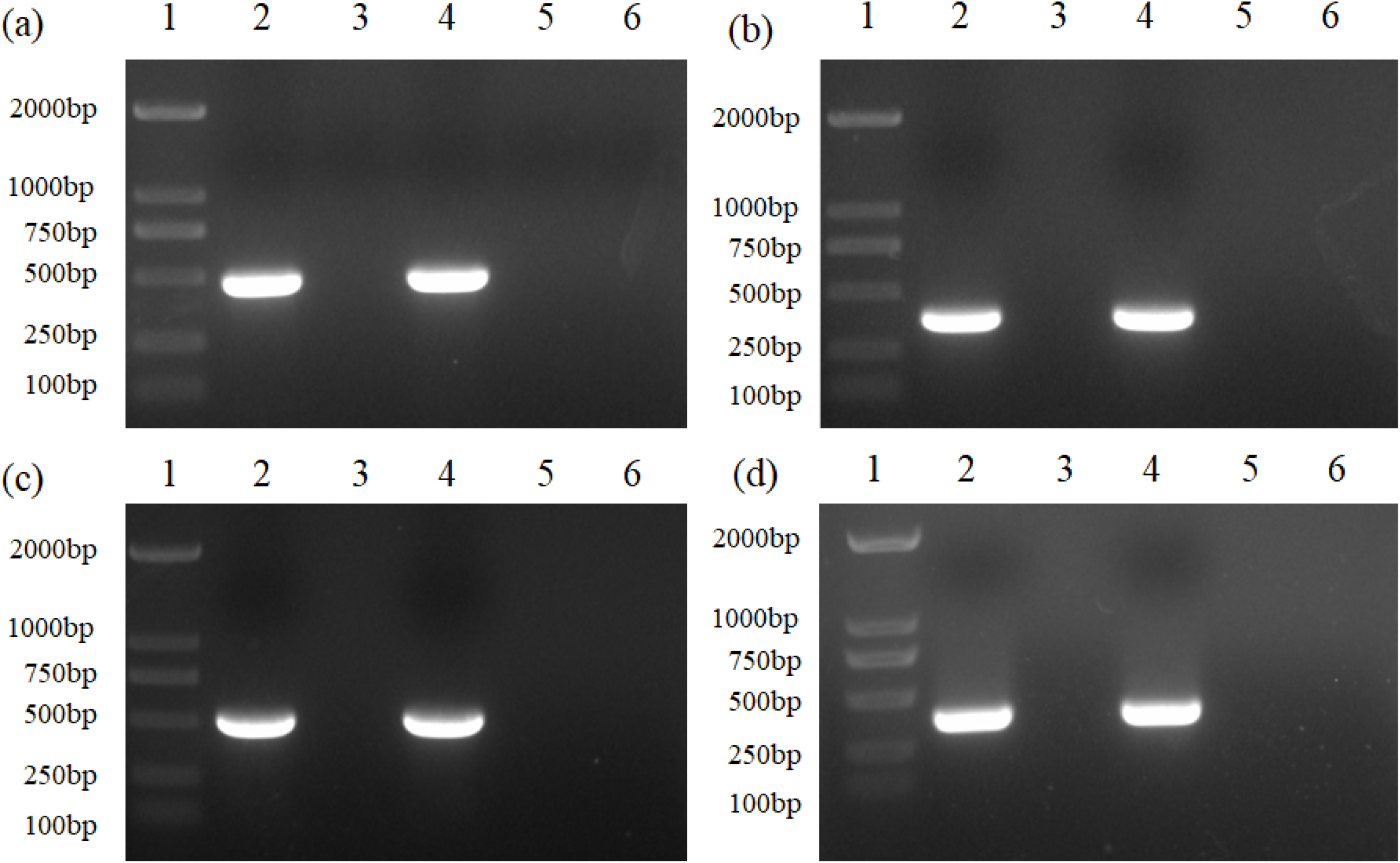
PCR verification of antibiotic resistance genes. (a). bl2d_oxa1; (b). aph33ib; (c). cml_e3; (d). tetb. Lane 1. marker 2000bp; lane 2. P4chr1; lane 3. M; lane 4. M+OMVs (single colony from TSA^+^ plate with antibiotic); lane 5. Modesto+OMVs(single colony from TSA^+^ plate without antibiotic plate); lane 6. negative control

The antimicrobial susceptibility test was performed with donor strain P4chr1, susceptible strain Modesto and 4 ARG transformed Modesto strains. The test result showed that the MIC values in these 4 strains did not increase compared with the antibiotic sensitive Modesto strain (Table 4), which may imply that the ARGs transferred by donor OMVs are not persistent in recipient cells after passaging, when the ARGs disappear or no longer express drug resistence. *A. paragallinarum* serovar A, B and C specific antisera were used in HA-HI test to measure serotypes of donor strain P4chr1 (serovar A), recipient strain Modesto (serovar C) and ARG transformed strains.

**Table 4.** Antibiotic susceptibility profiles of *A. paragallinarum* P4chr1, Modesto and OMVs transformed strains.

| Antibiotics | MIC (µg/ml) |  |  |  |  |  |
| --- | --- | --- | --- | --- | --- | --- |
|  | P4chr1 | M | M+OMVs<sup>tet</sup> | M+OMVs<sup>amp</sup> | M+OMVs<sup>chl</sup> | M+OMVs<sup>str</sup> |
| Tetracycline | 32 | 0.5 | 0.5 | 0.5 | 0.5 | 1 |
| Ampicillin | 128 | 1 | 1 | 1 | 1 | 1 |
| Chloramphenicol | 32 | 0.25 | 0.25 | 0.25 | 0.5 | 0.25 |
| Streptomycin | >512 | 2 | 2 | 2 | 1 | 2 |

**Table 5.** Typing results of *A. paragallinarum* P4chr1 and transformant serotype

| Antigen | Serum |  |  |  |  |  |  |  |  |
| --- | --- | --- | --- | --- | --- | --- | --- | --- | --- |
|  | Hp8(A) |  |  | 0222(B) |  |  | Modesto(C) |  |  |
|  | 800x | 1600x | 3200x | 800x | 1600x | 3200x | 800x | 1600x | 3200x |
| P4chr1 | ++++ | ++++ | +++- | — | — | — | — | — | — |
| Modesto | — | — | — | — | — | — | ++++ | ++++ | +++- |
| Modesto+OMVs<sup>tet</sup> | — | — | — | — | — | — | ++++ | ++++ | +++- |
| Modesto+OMVs<sup>amp</sup> | — | — | — | — | — | — | ++++ | ++++ | +++- |
| Modesto+OMVs<sup>chl</sup> | — | — | — | — | — | — | ++++ | ++++ | +++- |
| Modesto+OMVs<sup>str</sup> | — | — | — | — | — | — | ++++ | ++++ | +++- |

Note : “++++” 、 “+++—” 、 “++--”indicates 100,75,50 percents of hemagglutination-inhibition respectively;“—”indicates no hemagglutination-inhibition, Hp8, 0222 and Modesto are known serotype strains and are stored in the laboratory.

The HI result showed that both *A. paragallinarum* Modesto and transformants gave same titers with serovar C antiserum and no reaction with serovar A antiserum. The test assured that the transformants were derived from the recipient bacterium and they were clearly separated from serovar A donor strain P4chr1.

To further confirm this conclusion, we used the PCR-RFLP verification. The result showed that a 1.6kb fragment in hypervariable region of Hmtp210 was amplified for P4chr1, M and transforments. After digestion with restriction enzyme Bgl II, the PCR products were divided into two bands, 768 and 868 bp for serovar A and 1,284 and 339bp in the case of serovar C. The results showed that donor bacterium P4chr1 was type A, and both the recipient bacterium M and the transformants were type C (Fig 8).

**FIG 8.**
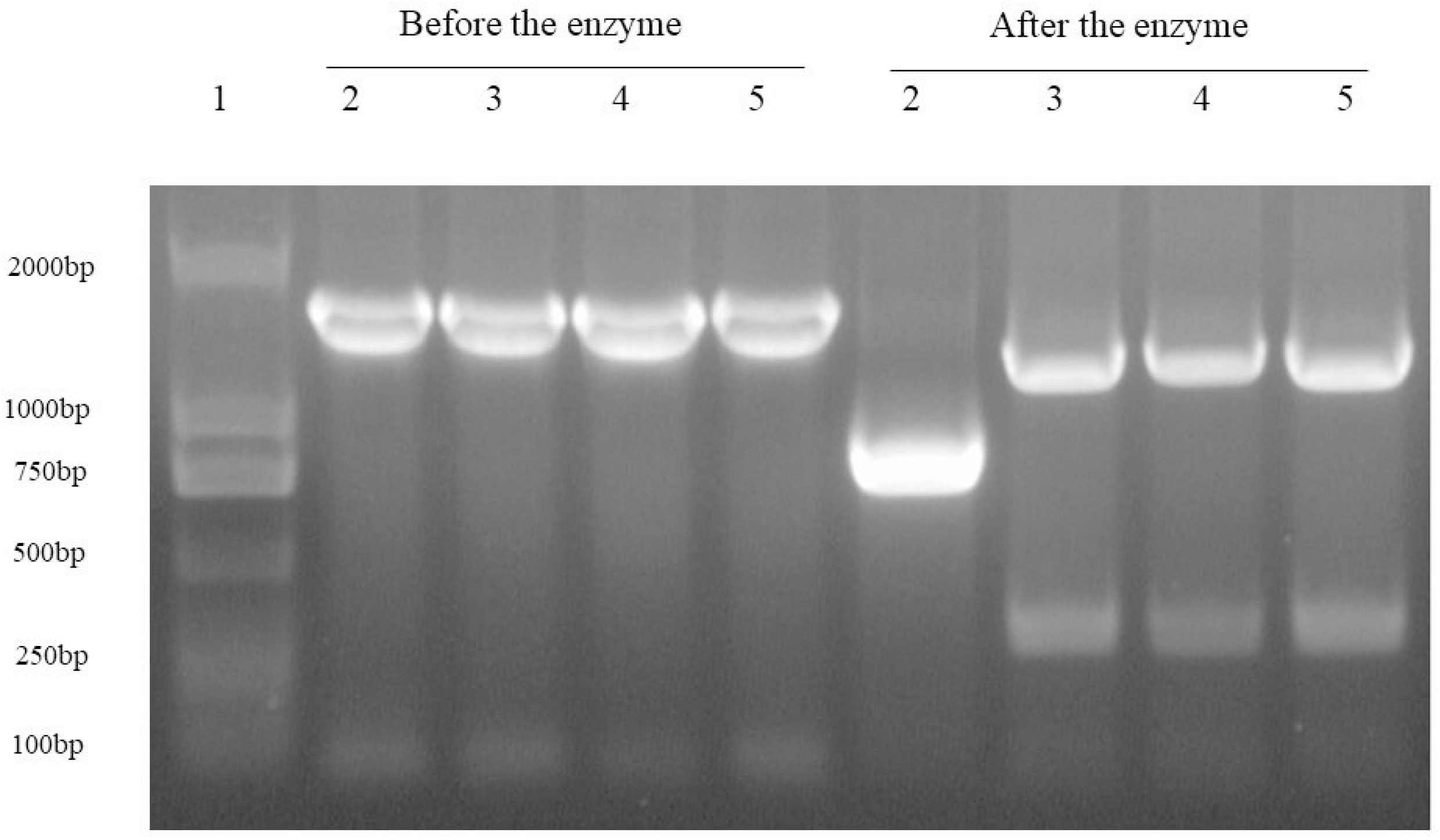
PCR-RFLP profile of the *A. paragallinarum* strains used in this study. Lane 1. marker 5000bp; lane 2. P4chr1; lane 3. M; lane 4. M+OMVs (single colony from TSA^+^ plate with antibiotic); lane 5. M+OMVs (single colony from TSA^+^ plate without antibiotic plate)

## DISCUSSION

Both pathogenic and non-pathogenic gram-negative bacilli secrete vesicles [9]. The fact that bacterial OMVs contain DNA (plasmid, chromosomal, and/or phage-associated) has been admitted [9, 18, 26]. Whether OMVs have complete genome as their parent cells do and if OMVs contain all the genetic information of the bacterial genome are unclear. By sequencing the purified *A. paragallinarum* OMVs, it was found that genome fragments contained in the OMVs are fragmented. In this study, we have sequenced multidrug-resistant *A. paragallinarum* strain P4chr1 and its OMVs, and performed comparative genomics analysis between the two genomes sequences (2.69 Mb and 2.77 Mb). The GC content, the number of coding genes and the metabolic pathways of the two genomes are very similar. The comparative genome circle map has to be magnified to see the gaps. Collinearity analysis found that each segment of the OMV genome sequence can be found in the corresponding part of the P4chr1 genome sequence. The whole genome orthologous gene cluster analysis revealed that the OMVs and P4chr1 had 2541 homologous cluster genes, with only a few differences, which may be caused by sequencing errors or fragmentation during OMV genome sequencing. These results indicated that the genome sequence of OMVs was derived from *A. paragallinarum* P4chr1, and the OMVs had almost all the genomic sequences of *A. paragallinarum* P4chr1, including virulence genes and antibiotic resistance genes. Within P4chr1 genome sequence, 11 ARGs are focused in 25kb region, forming a structure similar to tolerance island. After blast comparison, it is found that this sequence is very similar (99%) to a sequence of a field strain of *Pasteurella multocida* FCf83 from China, and the gene coding direction is also the same. These two bacteria belong to a same family as *Pasteurellaece*, which means that they have closer genetic affiliation and it may allow gene exchange in between easier. We don’t know whether this is the case, or if so, the sequence was transferred by transposition or insertion? In recent years, several important fuctions of OMVs have been revealed. the role of OMVs in horizontal transfer of antibiotic resistance genes among intra- and inter-species has been one of them [7]. In 2011, Carlos et al. reported the horizontal transfer of plasmids carrying the carbapenemase resistance gene OXA-24 in OMVs to *Acinetobacter baumannii* [13]. In 2015, Ho et al demostrated horizontal gene transfer mediated by *Porphyromonas gingivalis* OMVs. This bacterium (a fimA mutant) carried a 2.1 kb ermF-ermAM cassette in its fimA gene, encoding an erythromycin resistant gene. The cassette was transferred to the fimA gene of another *P. gingivalis* strain which is lack of such gene (erm gene) via OMVs isolated from the donor strain [16]. In 2019, Fulsundar et al proposed an optimized and detailed plan to test and confirm that OMVs-mediated ARGs can be transferred to *Acinetobacter baumannii* without plasmids in OMVs [12].

Here we have performed an experiment to test OMV-mediated HGT from *A. paragallinarum* P4chr1 strain who contains antibiotic resistance genes to antibiotic sensitive *A. paragallinarum* strain Modesto. As a result, we did succeed to see the ARG strain Modesto were survived on agar plates with ARGs, and its transformants passed several verification tests, i.e., ARG-PCR proved that the transformants amplifed corresponding ARG products, HA-HI test and PCR-RFLP confirmed that the transformants were derived from the recipient cells, not from donor cells. Howvers, the overall ARG transformation efficiency mediated by *A. paragallinarum* OMVs is low compaired with other reports which generally transfered ARGs carried in plasmid [13, 14, 17, 27]. In this HGT experiment, the highest transformation rate is 5.42% (Table 3) in streptomycin group and maybe there is higher copy number of streptomycin resistant gene in donor strain P4chr1. Another explaination says that each OMV contains merely a part of genomic fragments, some fragments are not containing intact ARGs, or not every vesicle contains DNA [27]. Tran and Boedicker considered that the ability to acquire DNA may depend on the species of the donor/recipient bacteria [28].

In our antimicrobial susceptibility test, the MIC values produced by transformant strains did not increase compared with antibiotic susceptible strain Modesto. Such unexpected outcome exposed a poor passaging ability of the transformants in this HGT study. It is proposed that after a gene is transferred into recipients, it must be integrated into chromosomal DNA in order to persist within the cells [16]. Ho et al presumed and proved that the erm gene they manipulated in the vesicles of the fimA mutant is flanked with fimA sequences at both ends, and that homologous DNA recombination occurs between vesicle donor DNA and the chromosomal DNA of the recipient [16].

In this study, there are no available homologous sequences flanked with drug-resistant genes in recipient bacterium Modesto (which can be confirmed by our genome sequencing results), so that the ARGs are unable to be incorporated into the chromosome of recipient cells through homologous recombination [16]. Moreover, even if there are homologous sequences, happening of gene recombination between vesicle donor DNA and recipient chromosomal DNA is just probable, and not as effective as most of HGT studies demostrated, where transferred genes were carried in plasmids [13, 14, 27].

Taken together, in this study we present complete genome sequencing data of *A. paragallinarum* P4chr1 and its OMVs and confirm that they are highly homologous. In addition, some drug resistance genes have been found in *A. paragallinarum* P4chr1 genome,and no drug resistance genes has been found in *A. paragallinarum* Modesto. Using purified OMVs from P4chr1 as vector, 4 AGRs are transfered into drug sensitive *A. paragallinarum* strain Modesto. Unfortunatelly, the ARG transformation efficiency and persistency are limited. Futher study is needed to understand OMV mediated HGT with chromosomal DNA based ARGs.

## Acknowledgements

The authors would like to thank Professor Xiao-ling Chen for her help.

## Author contributions

Hong-jun Wang designed the experiments. Jie Xu, Chen Mei, Yan Zhi, Zhi-xuan Liang and Xue Zhang performed the experiments. Jie Xu and Hong-jun Wang analyzed the results. Jie Xu and Hong-jun Wang wrote the paper. All authors read and approved the final manuscript.

